# A missense mutation (C667F) in β-dystroglycan results in reduced dystroglycan protein levels leading to myopathy and destabilization of the blood-brain and blood-retinal barrier protein network

**DOI:** 10.1101/2023.11.10.566559

**Authors:** Rui Lois Tan, Francesca Sciandra, Wolfgang Hübner, Manuela Bozzi, Jens Reimann, Susanne Schoch, Andrea Brancaccio, Sandra Blaess

**Affiliations:** Neurodevelopmental Genetics, Institute of Reconstructive Neurobiology, Medical Faculty, University of Bonn, 53127 Bonn, Germany; Institute of Chemical Sciences and Technologies “Giulio Natta” (SCITEC)-CNR, Rome, Italy; Biomolecular Photonics, Faculty of Physics, Bielefeld University, 33615 Bielefeld, Germany; Dipartimento di Scienze Biotecnologiche di Base, Cliniche Intensivologiche e Perioperatorie. Sezione di Biochimica. Università Cattolica del Sacro Cuore di Roma, 00168 Rome, Italy; Department of Neurology, Neuromuscular Diseases Section, University Hospital Bonn, 53127 Bonn, Germany; Synaptic Neuroscience Team, Institute of Neuropathology, Medical Faculty, University of Bonn, 53127 Bonn, Germany; School of Biochemistry, University Walk, University of Bristol, Bristol BS8 1TD, UK

## Abstract

Dystroglycan (DG) is a glycoprotein and extracellular matrix receptor consisting of an α-DG and a β-DG subunit encoded by the gene *DAG1*. A homozygous missense mutation (c.2006G>T), resulting in an amino acid substitution (p.Cys669Phe) in the extracellular domain of β-DG, causes severe Muscle-Eye-Brain disease with multicystic leukodystrophy. To investigate the mechanisms underlying the severe human pathology, we generated a mouse model of this primary dystroglycanopathy. We find that homozygous mutant mice show no obvious abnormalities during development and reach mature adulthood. However, α- and β-DG protein levels are significantly downregulated in muscle and brain of homozygous mutant mice. The mutant mice show a form of myopathy with late-onset and not fully penetrant histopathological changes in skeletal muscle and are impaired in their performance on an activity wheel. The brain and eyes of the homozygous mutant mice appear to be structurally normal, but the localization of mutant β-DG is altered in the glial perivascular endfeet (PVE) at the blood-brain- and blood-retina barrier resulting in a perturbed protein composition in the PVE. In conclusion, the mouse model of the C669F β-DG mutation does not seem to recapitulate the severe developmental phenotypes observed in human patients but represents a novel and highly valuable tool to study the impact of β-DG functional changes at the molecular level and to gain insight into the pathogenesis of primary dystroglycanopathies.

## Introduction

The dystroglycan (DG) protein complex is central in a number of physiological and pathological contexts, playing a particularly important role in skeletal muscle, brain and eye (Adams & Brancaccio, 2015). It is composed of two subunits, the extracellular and highly glycosylated α-DG and the transmembrane β-DG, which act as a molecular link forming an axis between the extracellular matrix and the cytoskeleton (Ibraghimov-Beskrovnaya *et al*, 1992; Ervasti & Campbell, 1993). The β-DG subunit establishes contacts with dystrophin and the cytoskeleton. DG is the major non-integrin cell-extracellular matrix adhesion complex and provides stability to various tissues such as skeletal and smooth muscle and the central and peripheral nervous systems. In addition, DG is involved in the stabilization of cell-matrix interfaces such as at the neuromuscular junction, the interface between endothelial cells and astrocytic endfeet at the blood-brain barrier, podocyte-glomeruli basement membrane contacts, and at the epithelia-connective tissue border in the lung (Barresi & Campbell, 2006; Bozzi *et al*, 2009; Winder, 2001). In addition, DG plays a very early and crucial role during mouse embryogenesis (Williamson *et al*, 1997). Mutations that disrupt the function of DG, its associated proteins or enzymes important for its post-translational maturation, lead to various forms of muscular dystrophy, which can be accompanied by distinctive eye, and brain phenotypes (Bianchini *et al*, 2014; Ackroyd *et al*, 2008; Liu *et al*, 2006).

Most of the DG-related muscular dystrophies result from alterations of the α-DG glycosylation shell observed in “secondary dystroglycanopathies” that are due to genetic abnormalities of glycosyltransferases involved in the glycosylation of α-DG (Muntoni *et al*, 2008; Fortunato *et al*, 2014; Endo, 2014). The N-terminal domain of α-DG (Brancaccio *et al*, 1997; Bozic *et al*, 2004) may also be involved, as it controls the binding of DG to glycosyltransferases or to additional factors important for the post-translational maturation of DG in the Golgi (Kanagawa *et al*, 2004). However, the exact relationship between the level of α-DG glycosylation and its pathophysiological consequences remains to be elucidated (Jimenez-Mallebrera *et al*, 2008). Although an increasing number of severe neuromuscular diseases resulting from *DAG1* mutations, the so-called “primary dystroglycanopathies”, have been identified (Hara *et al*, 2011; Geis *et al*, 2013; Özyilmaz *et al*, 2019; Dong *et al*, 2015; Dai *et al*, 2019), the underlying molecular basis of these diseases remains largely unknown (Brancaccio, 2019).

One of the primary dystroglycanopathies results in Muscle-eye-brain (MEB) disease with multicystic leukodystrophy, a severe neuromuscular condition with brain and eye abnormalities. This primary dystroglycanopathy arises from a homozygous missense mutation (c.2006G>T) resulting in an amino acid substitution (p.Cys669Phe) in the ectodomain of β-DG and has been described in two human patients (i.e. two sisters, aged 2 and 3 years) (Geis *et al*, 2013). Beyond the identification of the initial carrier family and the characterization of histomorphological features (analyzed at an early postnatal stage and with no follow-up available), our previous work in transfected cell lines showed that the β-DG C667F mutation in mice (murine topological counterpart to the human C669F mutation) leads to a defective trafficking process of the entire adhesion complex, such that the complex remains mostly engulfed in the endoplasmic reticulum (Signorino *et al*, 2018). No further biochemical or molecular data or follow-up of the two patients are available on this primary dystroglycanopathy, and it remains unclear how the mutation in β-DG leads to the destabilization of the whole DG complex (DGC) and to the severe symptoms observed in human patients.

To unravel the underlying mechanisms of the pathology in muscle, brain, and eyes, we generated and characterized a mouse model carrying the C667F mutation (corresponding to the C669F mutation in humans) within the ectodomain of β-DG. Our analysis shows that the mouse model of the C669F β-DG-associated MEB does not seem to recapitulate the developmental phenotypes observed in human patients, but that this mutation results in highly specific deficits in the glia-vascular unit in the central nervous system (CNS) and in late-onset histopathological changes in skeletal muscle. Taken together, our mouse model represents a novel and highly valuable tool to understand the mechanisms underlying certain aspects of the pathology of primary dystroglycanopathies and to study the impact of β-DG functional alterations at the molecular level.

## Material and Methods

### Animals

All mouse experiments were carried out with strict observance of protocols and guidelines approved by the University of Bonn Animal Care and Use Committee, Federal Government of Germany and European Union legislation. The protocols were approved by the Landesamt für Natur, Umwelt und Verbraucherschutz Nordrhein-Westfalen (Permit numbers: 81-02.04.2019.A415, 81-02.04.2019.A493). The mice were housed under controlled light (12:12 h light: dark cycle at an ambient temperature of 22°C). Water and mice chow were available *ad libitum*.

### Generation of mouse line

The *Dag1^C667F^* mouse line was developed by Genoway (Lyon, France) using a standard homologous recombination approach in embryonic stem cells. To generate the C667F mutation (corresponding to the C669F mutation in human) a point mutation (TGC>TTC at position 2698 in the cDNA sequence) was introduced into exon 5. The targeting vector containing the mutated exon 5 and flanking genomic regions (homology arms) was transfected into mouse ES cells. Neomycin (Neo)-resistant cells were screened by PCR and DNA sequencing. Positive clones were injected into blastocysts to generate chimeras. To excise the Neo cassette and generate the F1 generation (heterozygous point mutant knock-in mice) chimeras were crossed with mice expressing Cre in the germline. Pups from the F1 generation were then screened to test for germline transmission. Mice were generated on a C57BL6 background and then crossed onto a CD1 background. Heterozygous mice were crossed to obtain homozygous *Dag1^C667F/C667F^* mice.

### Genotyping

PCR amplification was performed to identify wildtype, heterozygous and homozygous mice. Genomic DNA was isolated from tail tips (neonates) or ear clips using sodium hydroxide digestion (Digestion with 75 μl NaOH for 1 h at 96°C followed by neutralization with pH 5.5 Tris HCl). PCR amplification was carried out using these conditions: 95°C for 30 s, 65°C for 30 s and 72°C for 30 s (30 cycles) with a final extension at 72°C for 8 min. The PCR products were loaded onto 1.5% agarose gel with ethidium bromide and photographed using a ChemiDoc and Image Lab software. Wildtype primers were: forward (F) 5’CCCCAGACTGGCCTTCAACTCATC3’; reverse (R): 5’AGTGCCCTATCACATGACATCCTGTCAC3’. Wildtype *Dag1* allele yields a 177 bp product, the *Dag1* C667F mutant allele a 268 bp product.

### Histology

#### Tissue preparation, brain, muscle, eye

Postnatal day (P) 0 (neonatal) and P7 (1 week) mice were decapitated; P60 (two months), P180 (six months) and 1-year-old mice (age range: 1.05 to 1.25 years) were euthanized using cervical dislocation. Eyes, brains, and whole hindlegs (for neonates) or hindleg muscles (for mice 1 week and older: upper and lower extensors, upper and lower flexors) were dissected. Eyes were fixed overnight in 4% paraformaldehyde (PFA), dehydrated in increasing concentrations of ethanol followed by xylene, embedded in paraffin and cut into 5 μm thick sections on a microtome. Brains were fixed in 4% PFA, cryoprotected in 15% and 30% sucrose and cryopreserved in OCT Tissue Tek (Sakura) on dry ice. P0 and P7 brains were cut into 14 μm thick sections onto adhesive microscope slides, while brains from P60 and older mice were cryosectioned at 40 μm thickness and collected as free-floating sections in anti-freeze solution (40% PBS, 30% ethylene glycol, 30% glycerol). Mouse muscle or hindlegs were snap frozen in isopentane chilled in liquid nitrogen and cryosectioned at 7 μm thickness. Note that for 1-year old mice, muscles, brain and eyes were isolated from animals that had access to a running wheel for 8 days before the tissues were collected.

#### Hematoxylin and Eosin staining

Paraffin sections of eye tissue were deparaffinized and rehydrated. Frozen sections were thawed and hydrated in PBS (brain tissue) or stained directly (muscle tissue). Sections were stained for 3 min in Hematoxylin, differentiated for 5 s (muscle) or 30 s (brain and eye) in 0.2% HCl 75% ethanol, blued for 5 min in running tap water, washed in double-distilled water for 1 min and stained with Eosin for 3-5 min. Sections were then dehydrated in increasing concentrations of ethanol followed by xylene and mounted with DPX (Merck) non-aqueous mounting medium.

### Immunostaining

For immunofluorescent staining of frozen sections, sections were re-fixed in 4% PFA for 10 min at room temperature (RT) and blocked in 10% NDS in PBS plus Triton X-100 (PBT) for 1 h at RT. For blocking and all the following steps 0.1% PBT was used for P0 and P7 brain tissue and muscle tissue at all stages while 0.3% PBT was used for adult brain tissue. Sections were incubated with primary antibody in 3% NDS PBT overnight at 4°C. To visualize brain endothelial cells, DyLight649 Lycopersicon Esculentum Lectin was added to the primary antibody solution (1:200; Vector Labs, Burlingame, CA, USA). Sections were washed 3 times for 5-10 min in PBT and incubated for 2 h at RT in secondary antibody and Hoechst (Abcam) in 3% NDS in PBT. Sections were washed 3 times for 5-10 min in PBT.

For immunofluorescent staining using the anti-β-Dystroglycan antibody (mouse, Novocastra), a “mouse on mouse” protocol was used. Sections were incubated in 10% NGS in 0.1% PBT for 30 min at RT. Sections were then incubated with unconjugated Fab fragments (Rockland) diluted at 1:100 in 1% BSA in 0.1% PBT for 1 h. Sections were washed 3 times for 5-10 min in 0.1% PBT followed by the standard immunofluorescent protocol described above but using DyLight549-conjugated goat-anti-mouse F(ab’)_2_ fragments as secondary antibody.

For immunofluorescent staining of paraffin sections, sections were deparaffinized and rehydrated. Antigen retrieval was done by boiling for 10 min in pH6.0 sodium citrate buffer in a pressure cooker. Sections were blocked for 1 h at RT with 10% NDS in 0.3% PBT and incubated overnight at RT with primary antibodies in 3% NDS in 0.3% PBT. To visualize brain endothelial cells, DyLight649 Lycopersicon Esculentum Lectin was added to the primary antibody solution (1:200; Vector Labs, Burlingame, CA, USA). Sections were washed 3 times for 5-10 min in 0.3% PBT and incubated with secondary antibodies and Hoechst (Abcam) in 3% NDS in 0.3% PBT for 1 h at RT. Sections were washed 3 times for 5-10 min in 0.1% PBT (P0 and P7 tissue) or in 0.3% PBT (adult tissue) and mounted with Aqua Polymount (Polysciences Inc.).

### Protein isolation and Western blot

Proteins were extracted from skeletal muscle (hindleg hip adductor and abductor, and thigh knee flexor complexes) and the whole brain of wildtype, heterozygous and homozygous mice (2 months, 6 months, 1 year) with phosphate buffer (PBS) containing 1% Triton-X100 (Sigma-Aldrich) and a Complete EDTA-free cocktail of protease inhibitors (Roche). Soluble proteins from skeletal muscle were incubated with Protein-A beads. The cleared fractions of skeletal muscle samples and total protein extracts of the brain were incubated with 500 μl of succinylated wheat germ agglutinin (WGA)–agarose beads (Vector Labs) at 4°C for 16 h. Beads were washed three times in washing buffer (WB, 1 ml PBS containing 0.1% Triton X-100 and protease inhibitors) and eluted with 300 μl WB containing 300 mM N-acetylglucosamine. 20 μg total protein extract or of WGA-enriched proteins were separated using 4-15% SDS–PAGE gels and were transferred to nitrocellulose membrane (Millipore, Bedford, MA, USA). Blots were probed with primary antibodies and then developed with horseradish peroxidase (HRP)-enhanced chemiluminescence (WesternBright ECL, Advansta, USA).

**Table 1.** List of antibodies.

| Antibody (species) | Source | Identifier | Titer/Volume/<br>Dilution (when applicable) |
| --- | --- | --- | --- |
| AKT (rabbit) | Fisher Scientific, Waltham, MA USA | RRID: 44609G | WB: 1:500 |
| α-Dystroglycan IIH6 (mouse) | Merck Millipore, Burlington, MA USA | RRID:AB_309828 | IHC: 1:100<br>WB: 1:1000 |
| α-Syntrophin (rabbit) | Alomone Labs, Jerusalem, Israel | RRID:AB_2756776 | IHC: 1:300 |
| Aquaporin 4 (rabbit) | Merck Millipore, Burlington, MA USA | RRID: AB_11210366 | IHC: 1:1000<br>WB: 1:1000 |
| BCL11A (mouse) | Abcam, Cambridge, UK | RRID: AB_2063996 | IHC: 1:1000 |
| β-Dystroglycan (mouse) | Novocastra, Newcastle upon Tyne, UK | RRID:AB_442043 | IHC: 1:200 |
| Calsequestrin (rabbit) | Thermo Fisher Scientific, Waltham, MA USA | RRID: AB_2071461 | IHC: 1:500 |
| CD31 (rabbit) | Abcam, Cambridge, UK | RRID: ab28364 | IHC: 1:50 |
| Dystroglycan (sheep) | R+D Systems, Minneapolis, MN USA | RRID: AB_10891298 | IHC: 1:30<br>WB: 1000 |
| Dystrophin (rabbit) | Abcam, Cambridge, UK | RRID: ab15277 | IHC: 1:200 |
| GFAP (chicken) | Merck Millipore, Burlington, MA USA | RRID: AB_177521 | IHC: 1:500 |
| GFAP (rabbit) | Dako, Santa Clara, CA, USA | RRID: AB_2811722 | IHC: Neat |
| Glutamine Synthetase(rabbit) | Merck Millipore, Burlington, MA USA | RRID: AB_2110656 | IHC: 1:500 |
| GLT-1 (guinea pig) | Merck Millipore, Burlington, MA USA | RRID:AB_90949 | IHC: 1:500 |
| KIR4.1 (rabbit) | Alomone Labs, Jerusalem, Israel<br>Thermo Fisher, Waltham, MA USA | RRID: AB_2040120<br>RRID: PA5-37137 | IHC: 1:200<br>WB: 1:500 |
| NeuN (mouse) | Merck Millipore, Burlington, MA USA | RRID: AB_2298772 | IHC: 1:500 |
| pAKT (rabbit) | Fisher Scientific, Waltham, MA USA | RRID: 44623G | WB: 1_500 |
| Pan-laminin (rabbit) | Abcam, Cambridge, UK | RRID:AB_298179 | IHC: 1:500 |
| PDGFRβ(mouse) | R&D Systems, Minneapolis, MN USA | RRID:AB_2162633 | IHC: 1:50 |
| S100β (rabbit) | Abcam, Cambridge, UK | RRID:AB_882426 | IHC: 1:500 |
| Tubulin-HRP (mouse) | Santa Cruz biotechnology, Dallas, TX US | RRID: Sc-23948 | WB: 1:1000 |
| Ubiquitin (rabbit) | Abcam, Cambridge, UK | RRID: AB134953 | WB: 1:200 |
| Alexa 488 donkey anti-mouse | Thermo Fisher Scientific, Waltham, MA USA | RRID:AB_141607 | IHC: 1:500 |
| Alexa 546 donkey anti-rabbit | Thermo Fisher Scientific, Waltham, MA USA | RRID:AB_2534016 | IHC: 1:500 |
| Alexa 488 donkey anti-chicken | Jackson ImmunoResearch, Ely, Cambridgeshire, UK | RRID:AB_2313596 | IHC: 1:500 |
| Alexa 488 donkey anti-sheep | Thermo Fisher Scientific, Waltham, MA USA | RRID:AB_141362 | IHC: 1:500 |
| Fab mouse IgG (H&L) antibody goat polyclonal | Rockland Immunochemicals, Pennsylvania, PA USA | RRID: AB_218897 | IHC blocking: 1:10 |
| Alexa 594 Affinipure Fab fragment goat anti-mouse IgG (H+L) | Jackson ImmunoResearch, Ely, Cambridgeshire, UK | RRID: AB_2338900 | IHC: 1:1000 |
| Anti-rabbit-HRP | Advansta, USA |  | WB: 1:10000 |
| Anti-sheep-HRP | R+D Systems, Minneapolis, MN USA |  | WB: 1:1000 |

### RNA isolation and quantitative RT-PCR

Total RNA was isolated from whole brain of 1-year-old wildtype, *Dag1^C667F/+^*or *Dag1^C667F/C667F^*mice using the RNeasy Mini Kit (Qiagen, Hilden, Germany) according to the manufacturer’s instructions. An additional on-column DNase treatment was performed to remove residual DNA. 1 μg of total RNA was reverse transcribed in a 20 μl reaction mix using the High-Capacity cDNA Reverse Transcription Kit (Applied Biosystems, Foster City, CA), following the manufacturer’s instructions. Quantitative RT-PCR was performed using a standard TaqMan® PCR protocol on a StepOne real time PCR System (Applied Biosystems) with primers specific for murine *Dag1*. The housekeeping gene hypoxanthine phosphoribosyltransferase (*Hprt*) was used as reference. The reactions were incubated at 50°C for 2 min and 95°C for 10 min, followed by 40 cycles of 95°C for 15 s and 60°C for 1 min. All reactions were run in triplicate.

### Imaging

Images of immunofluorescence-stained sections were acquired on an inverted Zeiss AxioObserver Z1 equipped with a Zeiss AxioCam MRm (Carl Zeiss, Oberkochen, DE). At 5x (EC PlnN 5x/0.16), 10x (EC PlnN 10x/0.3, Carl Zeiss, Oberkochen, DE) and 20X (EC PlnN 20x/0.5, Carl Zeiss, Oberkochen, DE) magnification, tile images were acquired with conventional epifluorescence. Tile images were stitched with Zen blue software (Zeiss, 2012).

At 63x (C-Apochromat, 63x/1.4 oil, Zeiss), images of immunofluorescence stainings were obtained at an inverted Zeiss AxioObserver equipped with a CSU-W1 Confocal scanner unit (50 mm pinhole disk, Yokogawa, Tokyo, JP). Z-stacks were acquired with laser lines 405 nm, 488 nm, 561 nm and 640 nm. Images taken are maximum intensity projections of these z-stacks.

Super-resolution 3D structured illumination microscopy was performed on an OMXv4 (GE Healthcare, Little Chalfont, UK) equipped with a 60x/N.A. 1.42 Olympus oil immersion lens and 4 separate 15-bit sCMOS cameras for fluorescent channel imaging. The corresponding laser excitation wavelengths were 405, 488, 568 and 642 nm. The raw images were processed with SoftWoRx 7.0.0 to construct 3D super resolution images. The 3D reconstruction was obtained with the molecular visualization software ChimeraX (Pettersen *et al*, 2021).

Brightfield images were visualized on a Zeiss Axio Scope.A1 microscope, acquired using AxioCam 503 Color and processed with Zen blue software (Zen lite, 2019).

### Voluntary wheel running

1-year-old mice (age range: 1.05 to 1.25 years; males n=4, females n=2) were individually housed in cages with an activity wheel (Scurry Mouse Misstep Wheel, Lafayette Instrument) for a period of 8 days. Mice were provided with free access to food and water, as well as nesting material and were allowed to run on the wheel at any intensity or duration. Mice had initially access to a 33-rung wheel (regular wheel) for 2 days. The complexity of the activity wheel was then increased according to the following scheme: mice were housed in a cage with an irregularly spaced 22-rung wheel (complex wheel) for 4 days and with an irregularly spaced 14-rung wheel (highly complex wheel) for 2 days. The running activity was recorded using Scurry Activity Monitoring Software (Lafayette Instrument), and through video recordings throughout the experimental period. Nocturnal (dark period in the light cycle) running activity was tallied and maximum speed, average speed, total duration per night, total distance per night were calculated using Microsoft Excel (version 16.16.27, Microsoft). Nocturnal running sessions, defined as a period of uninterrupted running on the wheel, were analyzed using Igor Pro software (version 8.04, Wavemetrics Inc., Oregon, USA) for average speed, maximum speed, average duration, average distance per session, as well as total number of sessions per night.

### Quantification

#### Tissue sections

Cross section areas and minimal Feret’s diameters of 6 months (males n=3, females n=2) and 1-year-old (age range: 1.05 to 1.25 years; males n=4, females n=2) mice quadriceps femoris muscle fibers were measured to quantify muscle fiber size variability as a surrogate for muscle fiber degeneration and regeneration. 10x Haematoxylin and Eosin light microscope images were traced and the cross-sectional area and minimal Feret’s diameter were quantified using ImageJ (Treat-NMD SOP DMD_M.1.2.001 Version 2.0). Percentage of muscle fibers with centralized myonuclei was also calculated for 6 months and 1-year-old mice to objectify muscle fiber regeneration. Muscle fibers with centralized myonuclei were manually counted using 10x Haematoxylin and Eosin light microscope images and expressed as a percentage of all muscle fibers in the image.

To quantify protein expression at the PVE, mean gray values of immunofluorescence-stained blood vessels (α- and β-DG, KIR4.1, AQP4 and α-Syntrophin) were measured in cerebral cortex and retina of 2-months-old (n=5) and 6-months-old (n=5) mice. Using 63x maximum intensity projections of z-stack images, blood vessels were manually traced based on Lectin staining, and the mean gray values of the traced areas were measured using ImageJ. Mean gray values for the marker of interest were normalized to the mean gray value of lectin.

#### Western blots

The densitometry analysis of bands was performed using Alliance Q9 Advanced UVITEC software and normalized to Tubulin.

#### Quantitative RT-PCR

The relative level for each gene was calculated using the 2^−ΔΔCT^ method(Livak & Schmittgen, 2001) and reported as fold change. Each experiment was repeated three times.

### Statistics

#### Western blots

One-way ANOVA followed by Sidak’s multiple comparison test was used for normalized densiometric measurement of control and mutant mice using GraphPad Prism 8.0 software (San Diego, California, USA). Error bars indicate SD.

#### Quantitative RT-PCR

One-way ANOVA followed by Sidak’s multiple comparison test was used on GraphPad Prsim 8.0 software (San Diego, California, USA).

*Histology and immunostaining:* Two-tailed Student’s t-test was used for the statistical analysis of mean gray values of blood vessels. One-way ANOVA with Sidak’s multiple comparison test was used for analysis of mouse weights on GraphPad Prism software 9.5. Histograms for muscle fiber cross section area and minimal Feret’s diameter were plotted. Variance coefficients (VC) of muscle fiber cross section area and minimal Feret’s diameter were calculated using the formula VC=1000 x standard deviation/mean (Briguet *et al*, 2004). VC were compared using one-way ANOVA with Sidak’s multiple comparison test. Statistical tests were performed in GraphPad Prism software 9.5.

*Voluntary wheel running*: Two-way ANOVA with Sidak’s multiple comparison test was used for analysis of maximum speed per night and per session, average speed per night and per session, total duration per night, total distance per night, average session duration, average session distance. Statistical tests were performed in GraphPad Prism software 9.5 and 10.0.3. Error bars indicate SEM or SD as specified in the Figure legends.

## Results

### Generation and characterization of the *Dag1^C66F7/C667F^* mouse line

To generate a mouse line carrying the C667F mutation (corresponding to the C669F mutation in humans) in the ectodomain of β-DG, a point mutation was introduced into exon 5 of *Dag1* (**Figure 1A**, see Materials and Methods for details). Mice heterozygous for the mutation (*Dag1^C667F/+^*) are viable, healthy, and fertile. Homozygous mice (*Dag1^C667F/C667F^*) survive to adulthood, are healthy and have no apparent behavioral phenotype in the home cage until over one year of age. Furthermore, homozygous mutants appear to be fertile (2 matings resulted in 2 litters of 2 and 4 pups, respectively). However, comparison with littermates showed that *Dag1^C667F/C667F^* males weigh significantly less than control littermates at 2 and 6 months of age, but not at 1 year of age (**Figure 1B**) We also found that *Dag1^C667F/C667F^* mice are not born with the expected Mendelian frequency (on average only 7.63 ± 1.98 % of mice per litter were homozygous), suggesting an embryonic phenotype that results in prenatal lethality but is only partially penetrant (**Figure 1C**).

**Figure 1.**
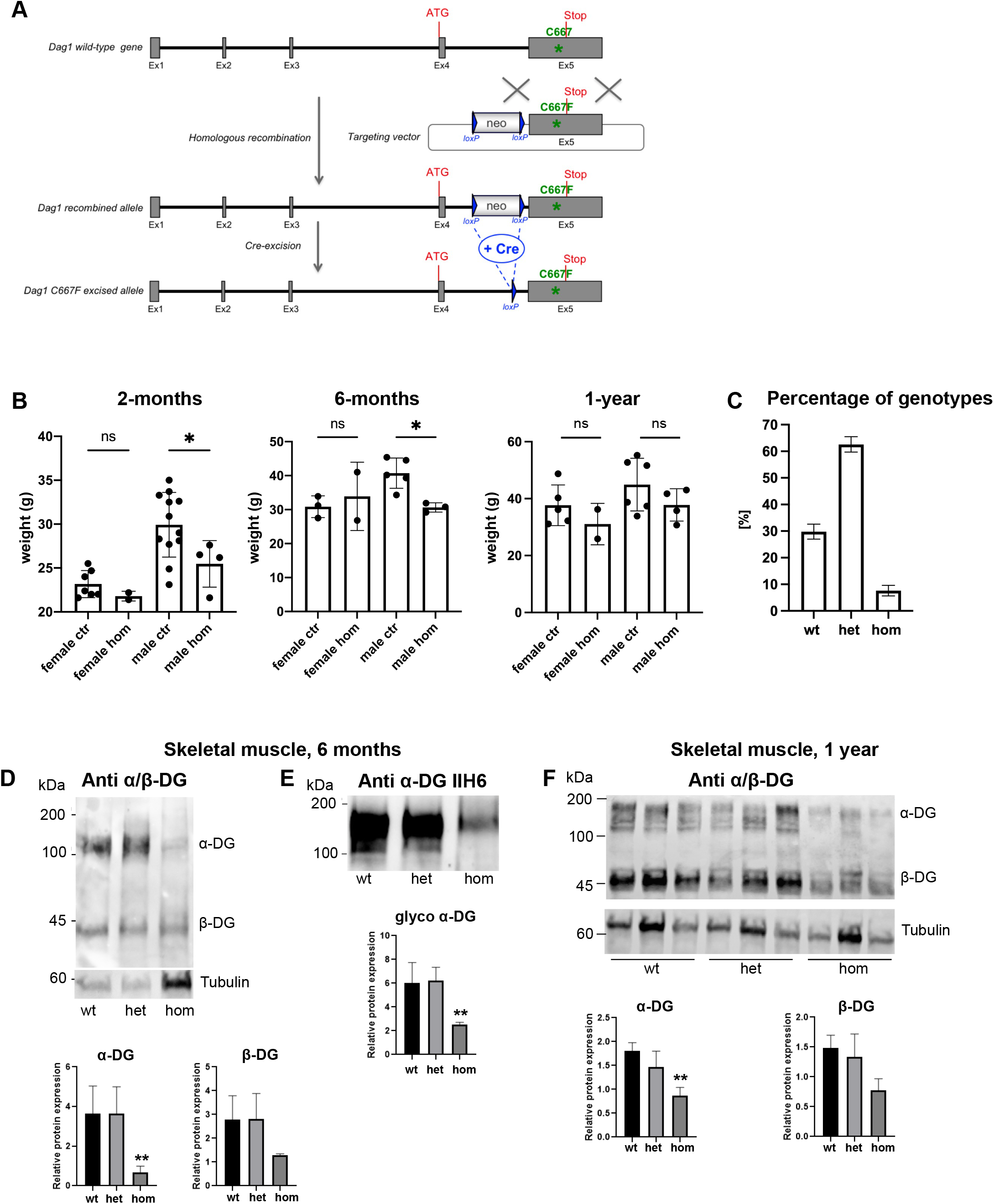
Reduced DG protein levels in skeletal muscle and brain tissue of *Dag1^C667F/C667F^ mice*. (A-C) Generation and characterization of the *Dag1^C667/C667^*mouse line. (A) Strategy for the development of a *Dag1^C667^* (human: C669F) point mutant mouse model: the model was developed using a standard homologous recombination approach in mouse embryonic stem (ES) cells. A targeting vector containing the mutated exon 5 (TGC>TTC) and flanking genomic regions (homology arms) was constructed and then transfected into ES cells. Neomycin (Neo)-resistant cells were screened by PCR and DNA sequencing. Positive clones were injected into blastocysts to generate chimeras. To excise the Neo cassette and generate the F1 generation (heterozygous point mutant knock-in mice), chimeras were crossed with Cre-expressing mice. Pups from the F1 generation were then screened to test for germline transmission. Shown are the wild-type allele, the targeting vector, the recombined locus found in ES cells and chimeras and the point mutant allele after in vivo excision of the neo cassette. (B) *Dag1^C667F/C667F^* (hom) male mice weigh significantly less than control (ctrl) mice (genotype: *Dag1^C667F/+^*or *Dag1^+/+^*) at 2 and 6 months of age, but not at 1 year of age. There is no significant difference between female control and mutant mice at the ages analyzed. (C) *Dag1^C667F/C667F^*mice are not born at the expected Mendelian frequency of 25%, indicating an embryonic phenotype that results in prenatal lethality but is only partially penetrant (47 litters, 361 mice). (D-F) Expression of DG was evaluated by Western blot performed on skeletal muscle tissues (hind limb hip adductor and abductor, and thigh knee flexor complexes) from wild-type (wt), heterozygous (het), and homozygous (hom) *Dag1*^C667F/C667F^ mice at 6 months or 1 year of age. Quantification of protein bands is reported as the average (6-month-old mice: n=4 for each genotype; 1-year-old mice: n=3 for each genotype) and presented as DG/Tubulin ratio. (D, F) α- and β-subunits were analyzed in WGA enrichments using a polyclonal α/β-DG polyclonal antibody recognizing both core proteins. (E) Glycosylated α-DG was detected using IIH6 monoclonal antibody. Error bars: (B, C) SEM, (D-F) SD. Statistical analysis was performed by one-way ANOVA with Sidak’s multiple comparison. *p<0.05; **p<0.01 (hom vs wt for D-F).

### DG protein levels are reduced in skeletal muscle and brain tissue of *Dag1^C667F/C667F^* mice

Next, we investigated whether the mutation leads to alterations in DG protein expression and processing in brain and muscle tissues. Based on our previous observations in transfected cell lines (Signorino *et al*, 2018) we expected an alteration in α/β-DG processing or trafficking and thus a potential reduction in protein levels in homozygous mice. Lysates and succinylated wheat germ agglutinin (WGA) enrichments from brain and muscle tissues were first subjected to a Western blot analysis with an antibody recognizing the core protein of both α- and β-DG. This showed that although DG was cleaved into its α- and β-subunits, α- and β-DG protein levels were substantially reduced in skeletal muscle and brain of *Dag1^C667F/C667F^* mice compared to wildtype or heterozygous controls (**Figures 1D,F**, **S1A**). We then used the anti-α-DG IIH6 antibody, which detects the glycan moiety responsible for α-DG binding to laminin and is commonly used to identify glycosylated α-DG. α-DG could be detected with this antibody in Western blot analysis of muscle and brain tissues from homozygous animals indicating that α-DG is glycosylated in *Dag1^C667F/C667F^* mice (**Figures 1E, S1B**). Thus, the C667F mutation results in reduced levels of DG but no alteration in its processing or glycosylation in the mouse model.

To investigate the mechanisms underlying the reduced DG levels we first performed quantitative RT-PCR of brain tissue to exclude that the reduced DG expression levels were due to reduced transcription of *Dag1*. This analysis revealed no significant difference in *Dag1* mRNA levels between homozygous, heterozygous, and wildtype animals (**Figure S1C**). In addition, analysis of polyubiquitination levels of total protein extracts from brain and muscle samples excluded hyperactivation of the proteasome-ubiquitin protein degradation pathway (**Figure S1D** and data not shown). Next, we investigated whether the reduced level of α-DG has an impact on signaling pathways downstream of DG-mediated cell-ECM interaction. In particular, we analyzed the pI3K/AKT pathway, and we found no differences in the ratio of pAKT/AKT between control and *Dag1^C667F/C667F^* mice (**Figure S1E**).

Given the reduced DG expression observed in Western blots of muscle and brain tissue from *Dag1^C667F/C667F^* mice, we next examined the effect of the mutation on DG protein localization in muscle fibers and on skeletal muscle integrity and function. In a further step, we performed a detailed analysis of DG localization and of potential phenotypes in the brain and retina of the homozygous mutant mice.

### α- and β-DG are localized at the sarcolemma in skeletal muscle of *Dag1^C667F/C667F^* mice

In muscle fibers, α- and β-DG are localized at the sarcolemma and α-DG binds to its extracellular binding partner, laminin. To determine whether the C667F mutation alters the localization of DG in muscle fibers, we first examined whether α- and β-DG could still be detected at the sarcolemma of muscle fibers in homozygous mice. To this end, we performed immunostaining for glycosylated α-DG, β-DG or α/β-DG core protein in conjunction with laminin on sections of the hindlimb muscle (tibialis, biceps, triceps) at multiple time points (**Figures 2A-F** and data not shown). For these and all subsequent experiments, both, heterozygous and wildtype mice were used as controls, as DG levels were not reduced in the heterozygous mice **(Figures 1, S1)**. Immunostaining revealed that α- and β-DG were present at the sarcolemma of homozygous mice and expression was not obviously reduced as compared to controls at any of the time points analyzed (**Figures 2B-H** and data not shown). Localization of laminin around the individual muscle fibers was also similar in mutants and controls (**Figures 2E, F**). Since the C667F mutant protein accumulates in the ER of transfected cell lines (Signorino *et al*, 2018), we investigated whether α- and β-DG protein might be partially retained in the sarcoplasmic reticulum (SR). To this end, we performed double staining for Calsequestrin (CASQ), a marker for the SR, and for either glycosylated α-DG or β-DG in muscle from 1-week and 6-months-old mice. We could not detect any obvious increase in α- or β-DG levels in the SR of the homozygous mutant mice at these stages (**Figures 2G, H** and data not shown). These data indicate that the mutation in the ectodomain of β-DG does not result in a severe disruption of DG processing or subcellular localization during muscle development and maintenance but rather in an overall reduction of DG protein levels that was only apparent in Western blot analysis.

**Figure 2.**
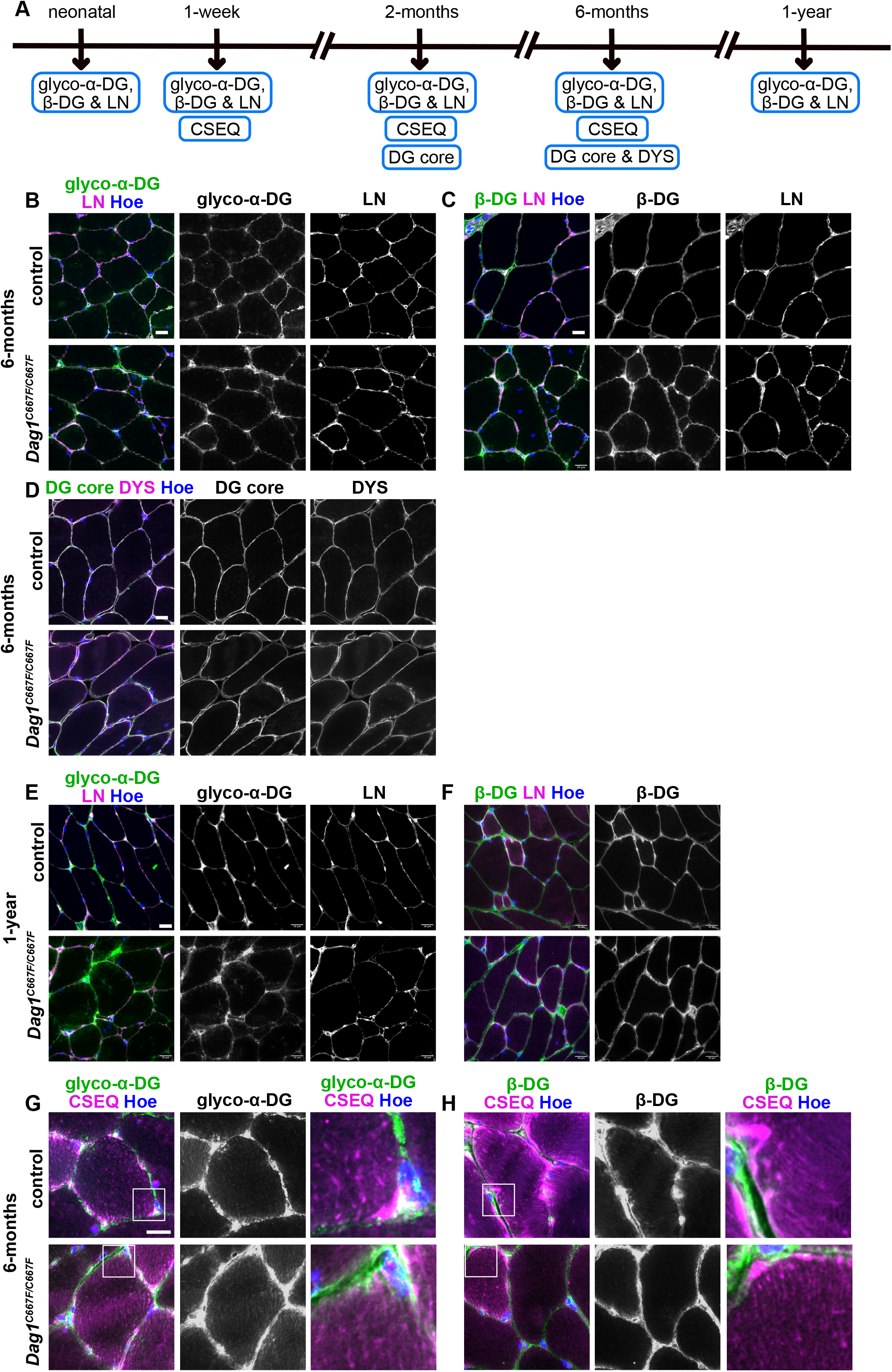
α- and β-DG are localized at the sarcolemma in skeletal muscle of *Dag1^C667F/C667F^ mice*. (A) Timeline summarizing which experiments were performed on which postnatal stages. (B-H) Cross-section through the quadriceps femoris muscle of 6-month- or 1-year-old mice. (B, C) Immunostaining for glycosylated (glyco) α-DG or β-DG in combination with laminin (LN) and Hoechst (Hoe) shows that DG is localized at the sarcolemma of the muscle fibers in control and *Dag1^C667F/C667F^*mice. (D) Immunostaining for the DG core protein (α/β-DG) and dystrophin (DYS). Both proteins are localized at the sarcolemma in control and mutant mice. (E, F) Immunostaining for glycosylated (glyco) α-DG or β-DG shows that DG is localized at the sarcolemma of muscle fibers in control and *Dag1^C667F/C667F^* mice at 1-year of age. (G, H) Immunostaining for glyco α-DG or β-DG in combination with the sarcoplasmic reticulum (SR) marker Calsequestrin (CSEQ) shows that DG is not trapped in the SR of *Dag1^C667F/C667F^*mice. The right panels are a higher magnification of the boxed area in the left panel. Scale bars: B-F: 20 μm, G, H: 10 μm.

### A late-onset histopathological phenotype is observed in skeletal muscle in a subset of *_Dag1_C667F/C667F* _mice_

Given the reduction in α- and β-DG protein levels in skeletal muscle of homozygous animals we examined whether the hindlimb muscle (m. Quadriceps femoris) of homozygous mutant mice had any histopathological changes (**Figures 3A-D** and data not shown). Both, heterozygous and wildtype mice were used as controls because the muscle histology of the heterozygous mice was indistinguishable from that of wildtype mice (data not shown). One sign of muscular dystrophy and the subsequent muscle fiber regeneration is the presence of nuclei in the center of the muscle fiber. Histological analysis of 2-months-old mice did not show an overt phenotype (**Figures 3B, C**). However, in 6-months and 1-year-old mice we found a substantial increase in the percentage of fibers with nuclei in the central region in two of the male homozygous mice (**Figure 3C**). Next, we performed a quantitative analysis of the muscle fiber cross-sectional area (2- and 6-months-old mice) and minimum Feret’s diameter (2-, 6-months and 1-year-old mice) in homozygous and control mice, since the degree of variability in muscle fiber size may be an indicator of muscular dystrophy (Briguet *et al*, 2004). This analysis showed no significant difference in the coefficient of variation (VC) of either the cross-sectional area or of the minimum Feret’s diameter of muscle fibers in homozygous mice compared to control mice at any of the analyzed stages (**Figure 3D** and data not shown). Even when these two measures were specifically compared between control males and males with an increased number of fibers with central nuclei, no significant difference could be detected (data not shown). Nevertheless, histograms of the minimum Feret’s diameter of mice at 6-months and 1-year of age showed a broadening and flattening of the distribution in male and female homozygous mutant mice compared to controls (**Figures S2A, B**). Taken together, these data suggest that the mutation in β-DG results in a subtle, late-onset and partially penetrant histopathological phenotype in the muscle of adult (older than 2-months) *Dag1^C667F/C667F^*mice.

**Figure 3.**
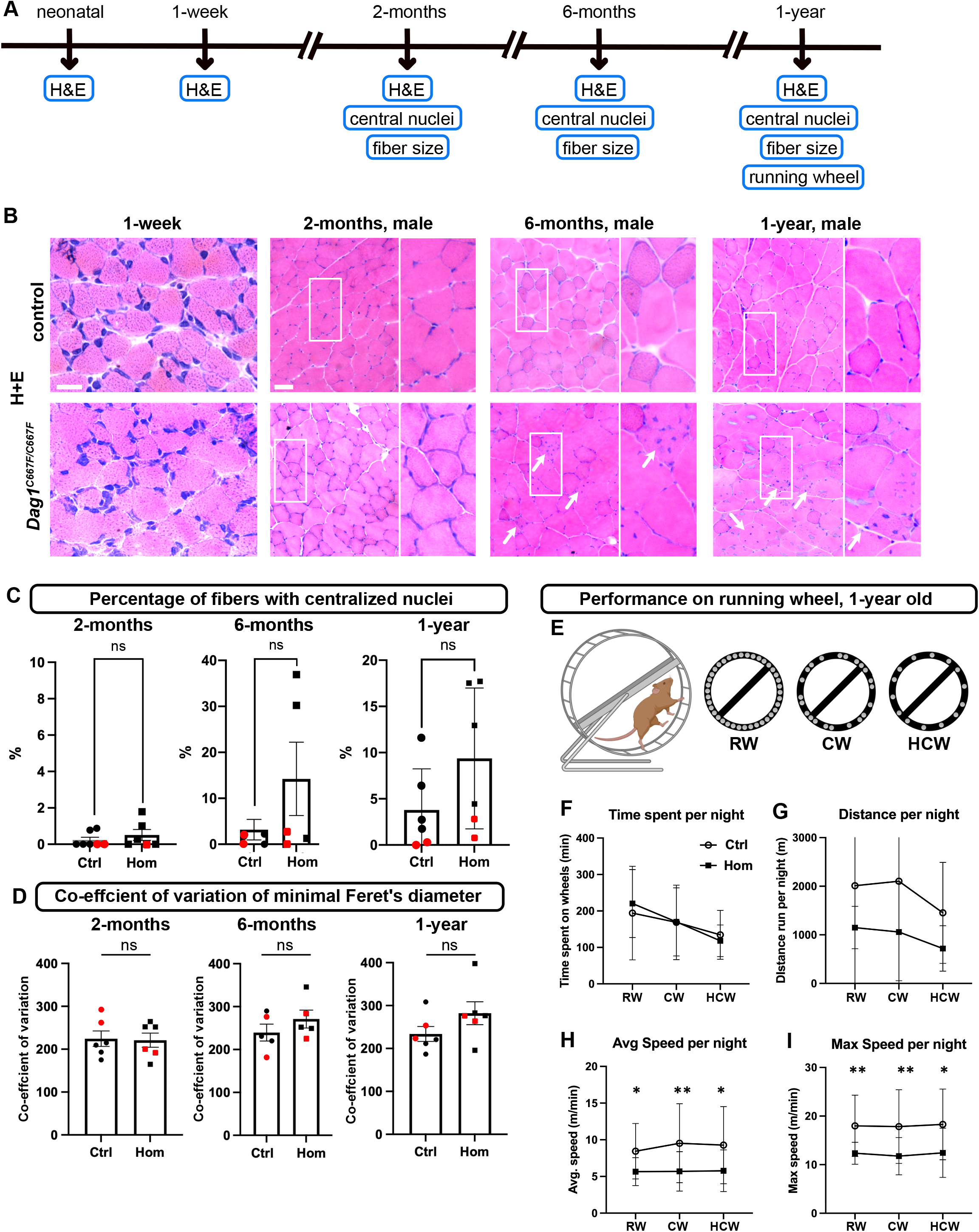
A late-onset pathological phenotype in skeletal muscle in *Dag1^C667F/C667F^* mice. (A) Timeline summarizing which experiments were performed at which postnatal stages. (B) Hematoxylin and Eosin (H&E) staining of cross-sections through the quadriceps femoris muscle at the indicated postnatal time points. 2- and 6-month- and 1-year-old muscle fibers: right panel is a higher magnification of the boxed area in the left panel. Arrows indicate the presence of centralized nuclei in 6-month- and 1-year-old muscle fibers. Scale bars: 1-week: 20 μm, 2-month- to 1-year-old: 100 μm. (C) Percentage of fibers with centralized nuclei. Note that while overall there is no significant difference between control (Ctrl) and *Dag1^C667F/C667F^*(Hom) mice, there are several male Hom mice (black squares) that have a particularly high percentage of fibers with centralized nuclei. (D) The variance coefficient (VC) of Feret’s diameter is not significantly different between Ctrl and Hom mice at the stages analyzed. (C, D) Black circles or squares: male mice, red circles or squares: female mice. Ctrl: control mice; Hom: *Dag1^C667F/C667F^* mice. Error bars indicate SEM. Statistical analysis was performed with unpaired t-test with Welch’s corrections (C, 2-month-old, D, 1-year-old), Mann-Whitney test (C, 6-month- and 1-year-old) or unpaired t-test (D, 2- and 6- month-old). (E-I) 1-year old mice were given access to an activity wheel in their home cage. (E) Mice had access to a 33-rung wheel (regular wheel, RW) for 2 days. Subsequently, the complexity of the wheel was increased: mice had access to an irregularly spaced 22-rung wheel (complex wheel, CW) for 4 days, followed by access to an irregularly spaced 14-rung wheel (highly complex wheel, HCW) for 2 days. Schematic created with Biorender. (F, G) Ctrl and Hom mice spent a similar amount of time per night on the RW, CW, or HCW, and the distance covered per night was not significantly different between Ctrl and Hom mice. (H, I) Average speed and maximum speed per night are significantly lower in Hom mice compared to Ctrl mice on RW, CW and HCW. Error bars: SEM. Statistical analysis was performed by two-way ANOVA with Sidak’s multiple comparison. *p<0.05; **p<0.01.

### Reduced running capacity in 1-year-old *Dag1^C667F/C667F^* mice

Given the reduced DG protein levels and the subtle histopathological phenotype in the hindlimb muscle of aged mice, we next investigated whether skeletal muscle function is altered in *Dag1^C667F/C667F^*mutant mice. To this end, 1-year-old control and *Dag1^C667F/C667F^*mutant mice were individually housed in cages containing an activity wheel and their performance was automatically monitored (Novak *et al*, 2012). Voluntary wheel running has been used in several studies to assess endurance in mice with impaired skeletal function (Elbaz *et al*, 2019; Mosca *et al*, 2013; Ruiz *et al*, 2022). In our paradigm, we also tested for effects on movement coordination: the wheel was made more complex over time by removing individual rungs (**Figure 3E**). Maximum running speed, average running speed, total running time and total running distance for individual sessions or the entire dark period were compared between control and *Dag1^C667F/C667F^* mutant mice for the three wheel types (**Figures 3F-I, S2C-H).** On the regular wheel, the average and the maximum running speed per dark phase and per session were reduced in the mutants as compared to the controls suggesting that their performance was impaired (**Figures 3H, I, S2F,G**). The performance levels observed on the regular wheel were maintained on the more complex wheel types in both control and mutant animals, indicating that motor coordination was not impaired in the *Dag1^C667F/C667F^*mutant mice (**Figures 3F-I, S2C-H**). The total time spent on the wheel per dark period, the length and number of individual sessions were not significantly different between control and mutant mice, indicating that endurance and motivation to run were not altered in the mutants (**Figures 3F, S2C,D).** Likely due to the decreased speed of the mutant mice, the total running distance per dark period appeared to be reduced (without reaching significance) compared to controls and the mean distance per session was significantly reduced for the complex and highly complex wheel (**Figures 3G, S2E)**. In conclusion, these data show that 1-year-old *Dag1^C667F/C667F^* mutant mice have a reduced running capacity as manifested by reduced speed compared to control mice, whereas time spent on the wheel, motivation and coordination are not overtly affected in the homozygous mutant mice, suggesting a muscle-specific phenotype.

### No obvious anatomical changes in brain or eye of *Dag1^C667F/C667F^* mice

Conditional inactivation of *Dag1* in the mouse brain results in a detachment of radial glia endfeet from the basement membrane at the pial surface of the brain and a collapse of radial glia fibers. This results in the disruption and partial loss of the basement membrane and layer defects in the cerebral cortex and cerebellum, and fusion of the cerebellar folia (Moore *et al*, 2002). Thus, if this mutation in β-DG affects DG expression levels, localization or function in the developing brain, homozygous mutant mice may have anatomically altered cortical structures. However, analysis of the layers in the cerebral and cerebellar cortex using Hoechst or neuronal markers (BCL11A for cortical neurons) at different postnatal time points (**Figure 4A**) did not reveal any changes in the organization of the cerebral and cerebellar cortex (**Figures 4B-E** and data not shown). Furthermore, the distribution of GFAP-positive astrocytes and the localization of laminin at the pial surface appeared to be normal in the homozygous mutants (**Figures 4C-E** and data not shown).

**Figure 4.**
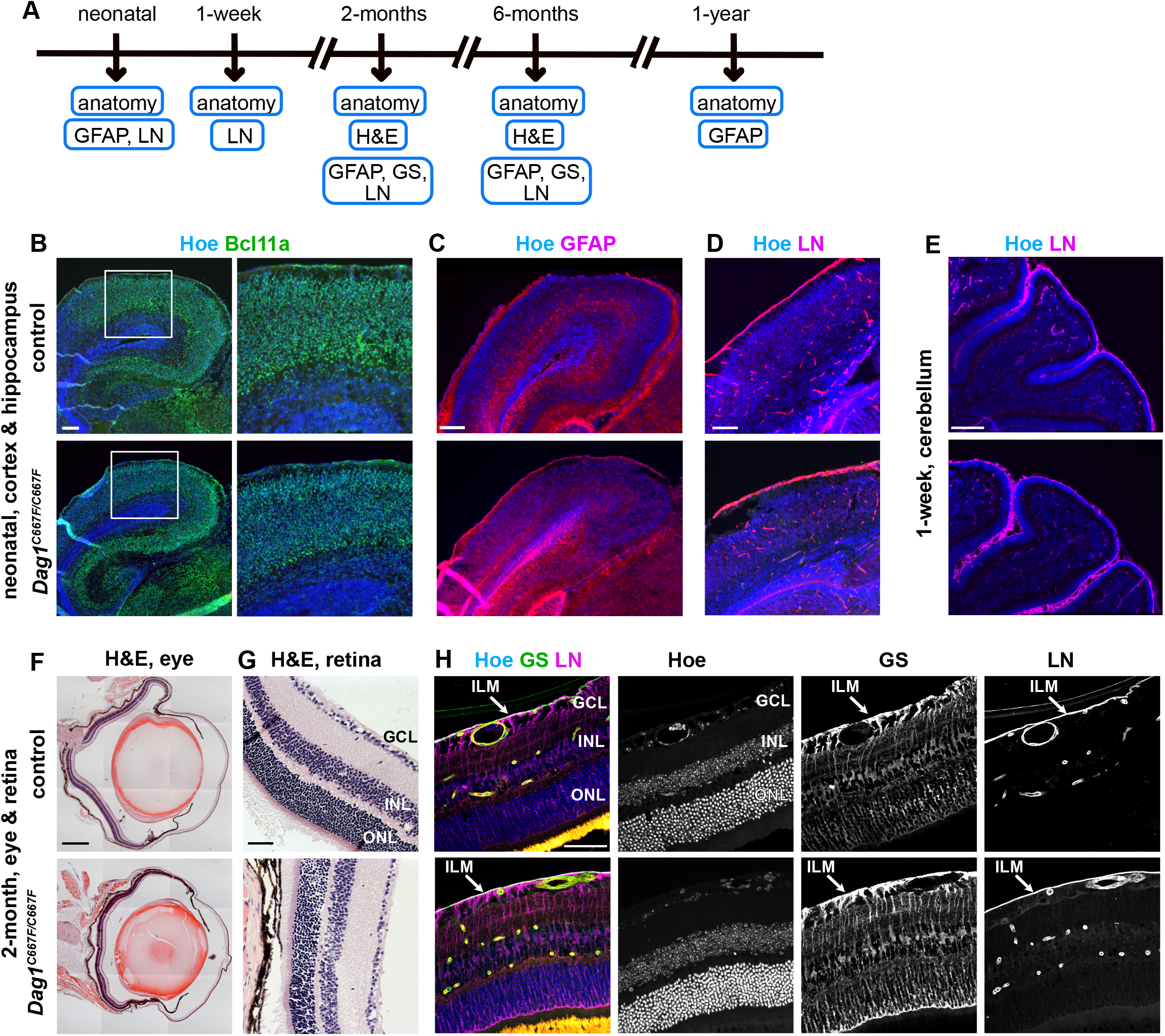
Brain and eye show no obvious anatomical changes in *Dag1^C667F/C667F^* mice. (A) Timeline summarizing which experiments were performed at which postnatal stages. (B-E) Analysis of brain anatomy in neonatal control and *Dag1^C667F/C667F^*mice. (B, C) Immunostaining for the cortical neuron marker BCL11A (B) or the astrocyte marker GFAP (C) in combination with Hoechst (Hoe) shows that the cortex, cortical layers and astrocyte distribution are not altered in neonatal *Dag1^C667F/C667F^*mice. (D) Laminin (LN) is expressed around blood vessels and at the pial surface of the neonatal in control and *Dag1^C667F/C667F^* mice. (E) Cerebellar folia and the pial basement membrane (LN positive) covering cerebellar fissures and folia are established in the mutant mice. (F-H) Analysis of eye and retina phenotype in 2-month-old control and *Dag1^C667F/C667F^* mice. (F, G) Hematoxylin and Eosin (H&E) staining of cross-sections through the eye (F) and retina (G) shows no obvious structural defect in the *Dag1^C667F/C667F^*mice. GCL: ganglion cell layer; INL: inner nuclear layer; ONL: outer nuclear layer. (H) Immunostaining for LN and glutamine synthetase (GS), a marker for Müller glia cells, in combination with Hoechst shows that the anatomical structure of the retina is not altered in the *Dag1^C667F/C667F^* mice. The inner limiting membrane (ILM) is present in homozygous mutant mice, as indicated by the presence of LN and GS-positive end feet of Müller glial cells at the ILM. Scale bars: B-E: 200 μm, F: 500 μm, G, H: 50 μm.

In addition to severe anatomical brain abnormalities, MEB disease associated with the DG C669F mutation is characterized by defects in the visual system such as congenital glaucoma, myopia, retinal atrophy and/or juvenile cataracts in the human patients (Geis *et al*, 2013). To investigate whether these phenotypes are recapitulated in our mouse model, we isolated eyes at different postnatal and adult stages (**Figure 4A**). Histological analysis of the eyes did not reveal any overt phenotypes, except for one mouse at two months of age with cataract in one eye (1 out of 12 eyes). In this eye, the ganglion cell layer was thinner and disorganized, which is also typically observed in glaucoma which presents with a loss of retinal ganglion cells (Munemasa & Kitaoka, 2013). In the developing retina, DG is required to maintain the structural integrity of the inner limiting membrane (ILM) (Clements *et al*, 2017). Loss of DG function results in ILM degeneration and deficits in migration and axon guidance, as well as altered anatomical arrangement of neurons and retinal thinning. Histological examination of the retina in the *Dag1^C667F/C667F^* mutants did not reveal any obvious changes in retinal organization (**Figures 4G, H** and data not shown). Immunostaining for glutamine synthetase (GS), a marker for Müller glia cells, which span almost the entire retina, showed that these cells were structurally normal. Finally, immunostaining for laminin showed that the ILM was properly established in the homozygous mice (**Figure 4H**, **Figure 7B** **and data not shown**).

Taken together, these data indicate that the mutation in β-DG does not disrupt the function of DG in organizing basement membranes at the pial surface of the brain or in the ILM in the retina in *Dag1^C667F/C667F^*mice. In addition, radial glia and Müller glia cells appear to be established and maintained and the anatomical development of the mouse brain and the retina are not overtly affected in the homozygous mutant mice.

### The molecular composition of the glia-vascular unit is impaired in the brain of *Dag1^C667F/C667F^ mice*

It has previously been shown that complete inactivation of *Dag1* in the CNS leads to alterations in the perivascular endfeet (PVE) of astrocytes and the blood-brain barrier. PVE are highly specialized astrocytic processes that envelop the brain’s vasculature (**Figure 5B**). α- and β-DG are expressed in the PVE, but also in endothelial cells (Zaccaria *et al*, 2001; Menezes *et al*, 2014).

**Figure 5.**
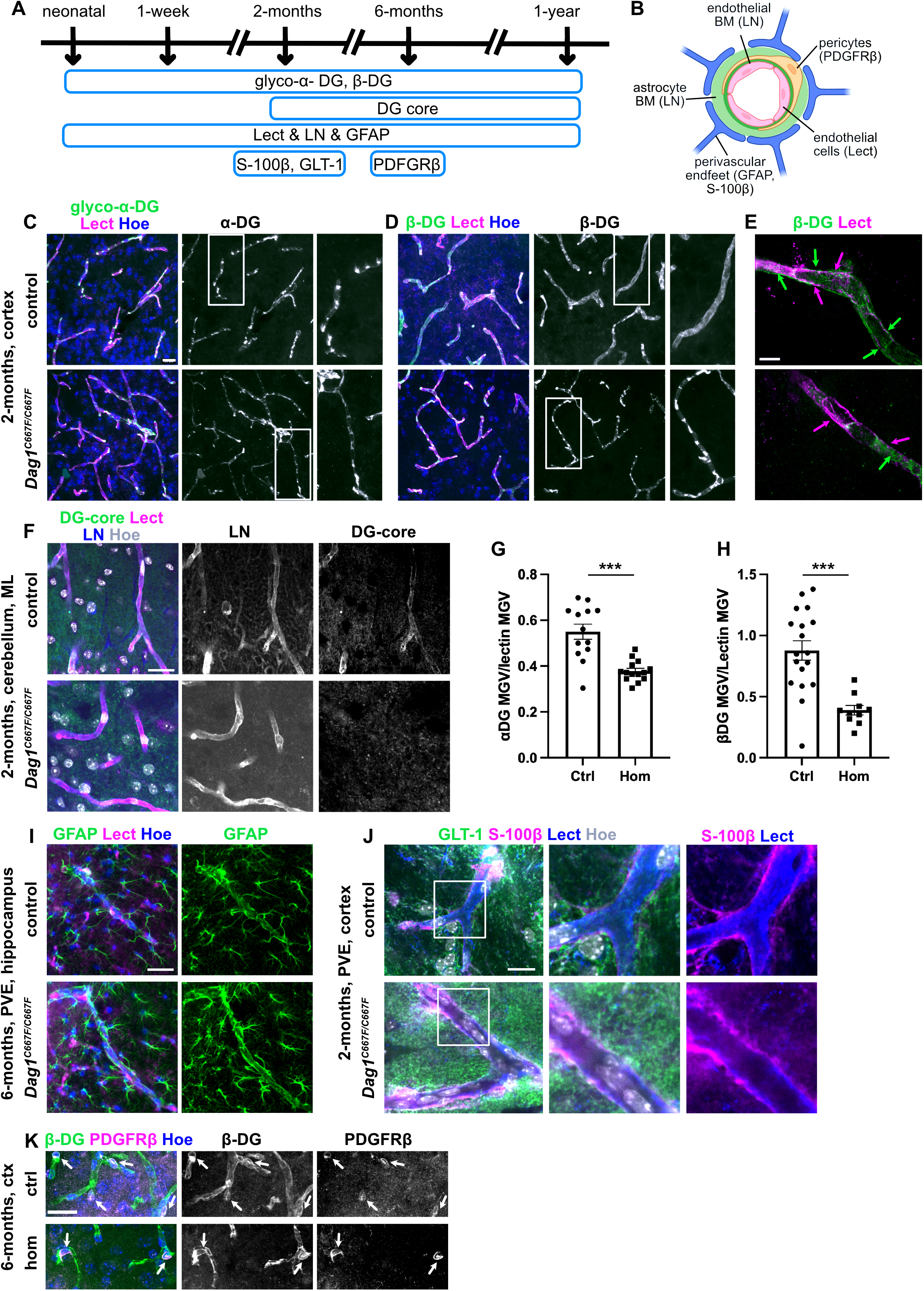
Perivascular endfeet are formed in the brain of *Dag1^C667F/C667F^* mice. (A) Timeline summarizing which experiments were performed at which postnatal stages. (B) Schematic representation of the cellular and ECM components of the blood-brain barrier. In brackets: markers expressed in the different components. BM: basement membrane. LN: Laminin, Lect: Lectin. Diagram generated with Biorender. (C-E) Immunostaining for glycosylated (glyco) α-DG or β-DG in combination with lectin (Lect) to visualize blood vessels and Hoechst (Hoe) in the cortex of 2-month-old mice. α-DG and β-DG are localized at blood vessels in *Dag1^C667F/C667F^* mice, but β-DG is abnormally clustered. (E) Super-resolution microscopy of a single blood vessel (Lectin-positive, magenta arrows) in control or homozygous mutant mice showing that β-DG (green arrows) is present in the perivascular space in control but not in *Dag1^C667F/C667F^* mice. 3 µm thick maximum intensity projections of z-stacks are shown. (F) Immunostaining for the α/β-DG core protein (DG-core) and laminin (LN) in combination with Lect and Hoe in the molecular layer of the cerebellum of 2-month-old mice. α/β-DG cannot be detected in the cerebellum of the homozygous mutant mice, whereas LN is still present. (G, H) Quantification of immunofluorescence intensity (mean gray value, MGV) of α-DG or β-DG expression in the PVE normalized to the MGV of Lectin shows that expression of both proteins is reduced in the PVE of the cortex in 2-month-old homozygous mutant animals compared to controls. n=5 Ctrl and 5 Hom animals, MGV was analyzed in 3 images per animal and included in the statistical analysis. Error bars: SEM. Statistical analysis was performed by unpaired t-test. ***p<0.001. (I) GFAP immunostaining in combination with Lect and Hoe visualizing astrocytes and their PVE surrounding a blood vessel in the hippocampus of 6-month-old control and *Dag1^C667F/C667F^* mice. (J) Immunostaining for GLT-1 to visualize astrocytes and S-100 β, which localizes to PVE, shows that PVE are still present in the cortex of homozygous mutant mice. (K) Immunostaining for β-DG and PDGRRβ to label pericytes. Pericytes (arrows) are located in the perivascular space and express β-DG in the cortex (ctx) of control (crtl) and *Dag1^C667F/C667F^* (hom) mice. Scale bars: C, D, E, F, I, K: 20 μm, E: 5 μm, J: 10 μm.

Immunostaining for glycosylated α-DG or β-DG in combination with lectin to label blood vessels in the cerebral cortex showed that α- and β-DG were localized at blood vessels at the analyzed stages in control and mutant brains (**Figures 5A, C, D**, **S3A,B** and data not shown). However, the intensity of the fluorescent signal for either α-DG or β-DG at the PVE appeared to be reduced in 2-months-old homozygous mutant animals compared to controls (**Figures 5G, H** and data not shown), and β-DG was abnormally clustered around blood vessels in the brain of *Dag1^C667F/C667F^* mice aged 2 months or older (**Figure 5D** and data not shown). Super-resolution microscopy of lectin and β-DG revealed that the perivascular localization of β-DG (surrounding the lectin staining on the extravascular site) was lost in homozygous mutant animals (**Figure 5E**, **Movies S1 and S2**). Furthermore, the DG core protein could be detected in the PVE in the molecular layer of the cerebellum in control mice, whereas it was absent in the PVE of *Dag1^C667F/C667F^* mice (**Figure 5F**). Labeling with GFAP, GLT-1 (Glutamate transporter 1, also known as EAAT1), and S-100β, three proteins expressed in astrocytes, two of which (GFAP, S-100-β) are known to localize to the PVE, showed that the PVE are formed in *Dag1^C667F/C667F^* mice (Langer *et al*, 2017) (**Figures 5I,J, S3C** and data not shown). Laminin, a component of the basement membrane between blood vessels and the PVE, was preserved in the homozygous mutant animals (**Figure 5F**) as were the PDGFRβ-positive pericytes in the perivascular space (**Figure 5K**) (Thomsen *et al*, 2017). Interestingly, the co-immunostaining for β-DG and PDGFRβ showed that β-DG is expressed in pericytes. In conclusion, these data suggest that the reduced presence and clustering of DG along blood vessels is likely a consequence of DG mislocalization in the PVE rather than structural changes in the PVE.

To investigate whether the altered localization of DG results in aberrant localization of other proteins important for blood-brain barrier function (**Figure 6B**), we examined the expression of Aquaporin 4 (AQP4), which is highly enriched in the PVE and plays a critical role in brain water homeostasis, at multiple time points (**Figures 6A, C-E** and data not shown). AQP4 was no longer properly localized to the PVE in homozygous mutant animals and could only be detected at very low levels around some blood vessels (**Figures 6C, D**). Western blot analysis of brain tissue showed that the absence of AQO4 in PVE was not caused by an overall reduction in protein levels (**Figure 6E**), suggesting that this phenotype is due to mislocalization of the water channel rather than reduced expression levels. AQP4 is anchored to the PVE membrane by components of the DGC complex (Cherkaoui *et al*, 2021; Nicchia *et al*, 2008; Sato *et al*, 2018; Lien *et al*, 2012). Therefore, we next examined the localization of dystrophin (DYS) and α-syntrophin (α-SNT) at the PVE. Immunostaining showed that both proteins were no longer localized to the PVE in the *Dag1^C667F/C667F^* mice at 2-months of age or older suggesting that the C667F mutation results in altered interactions of β-DG with the intracellular components of the DGC (**Figures 6F-H** and data not shown). Finally, we examined the localization of the inwardly rectifying K^+^ channel KIR4.1 (also known as KCNJ10), another important component of the blood-brain barrier that is also anchored to the PVE membrane by the DGC (Jukkola & Gu, 2015; Fujimoto *et al*, 2023). We found that the localization of KIR4.1 to the PVE appeared to be reduced in homozygous mutant mice as compared to controls, whereas the overall protein levels of KIR4.1 were not reduced in mutant animals (**Figures 6I-K**). Interestingly, AQP4, α-SNT and KIR4.1 were still localized around blood vessels in the brain of 1-week-old homozygous animals (**Figures S3D-F**), suggesting that DG is important for maintaining the molecular composition of the PVE during the blood-brain barrier maturation. In conclusion, these data indicate that the *Dag1^C667F^* mutation results in a specific impairment of DG function in the molecular organization of the PVE in the mature mouse brain.

**Figure 6.**
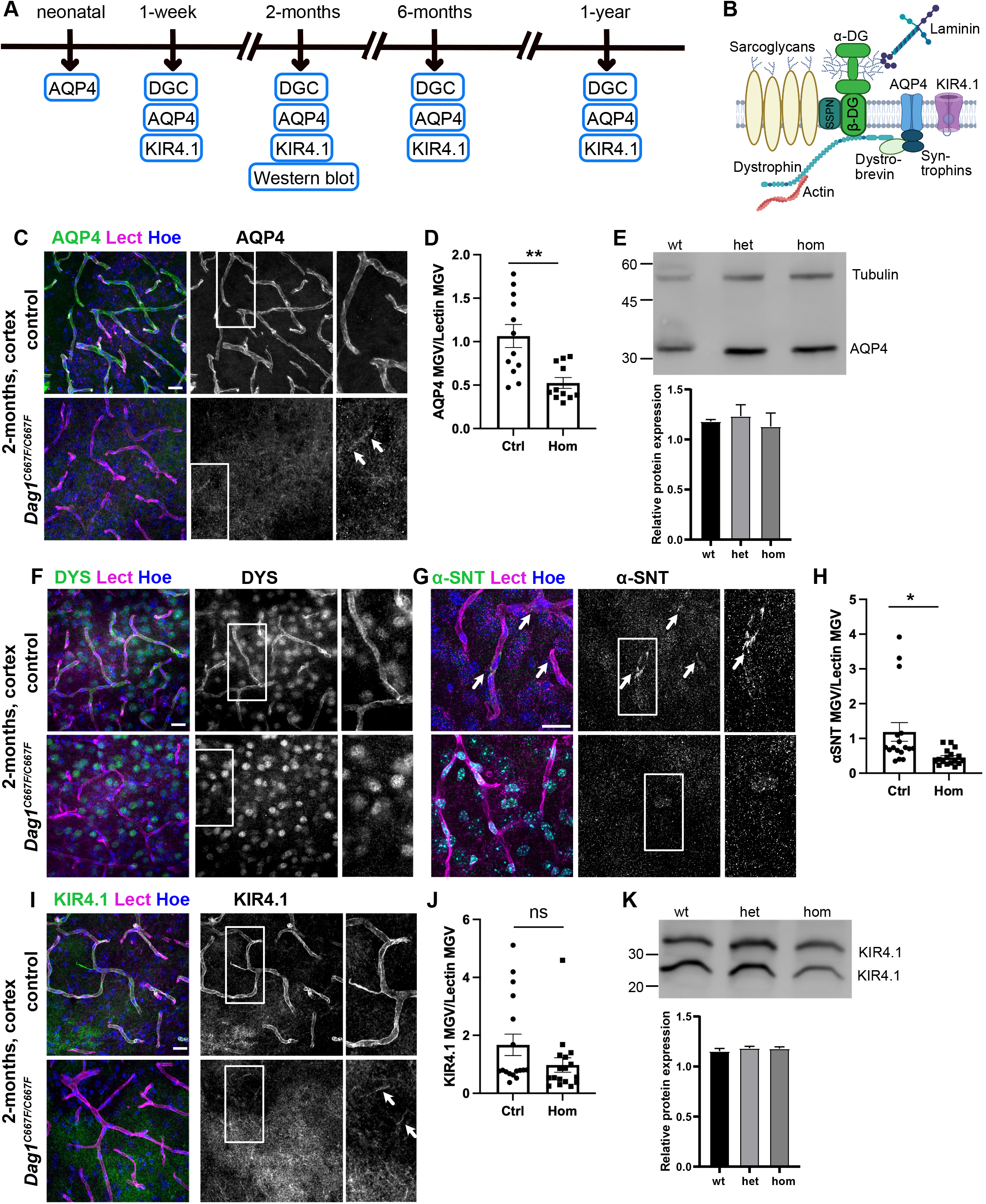
The molecular composition of the glia-vascular unit is disrupted in the brain of *Dag1^C667F/C667F^ mice*. (A) Timeline summarizing which experiments were performed on which postnatal stages. (B) Schematic representation of the DGC at the blood-brain barrier SSPN: Sarcospan, AQP4: Aquaporin 4, KIR4.1: inwardly rectifying K^+^ channel. Schematic created with Biorender. (C) Immunostaining for AQP4 in combination with Lectin and Hoechst shows that AQP4 is almost completely absent from the perivascular endfeet (PVE) in the cortex of 2-months old *Dag1^C667F/C667F^* mice. (D) Quantification of immunofluorescence intensity (mean gray value, MGV) of AQP4 in PVEs of 2-month-old control (Ctrl) and *Dag1^C667F/C667F^* (Hom) mice. The MGV of AQP4 is normalized to the MGV of Lectin. (E) Western blot for AQP4 showing that overall protein levels are not reduced in the brain of 2-month-old homozygous (hom) *Dag1^C667F/C667F^* mice compared to heterozygous (het) or wild-type (wt) mice. (F, G) Immunostaining for Dystrophin (DYS) and α-Syntrophin (α-SNT) in combination with Lectin and Hoechst. DYS and α-SNT are almost completely absent from the PVE in the cortex of 2-month-old homozygous mutant mice. (H) Quantification of immunofluorescence intensity of α-SNT in PVEs of 2-months old control and *Dag1^C667F/C667F^*mice. The MGV of α-SNT is normalized to the MGV of Lectin. (I) Immunostaining for KIR4.1 in combination with Lectin and Hoechst. KIR4.1 expression appears to be reduced in the PVE in the cortex of 2-month-old homozygous mutant mice. (J) Quantification of KIR4.1 immunofluorescence intensity in PVE of 2-month-old control and *Dag1^C667F/C667F^* mice. The MGV of KIR4.1 is normalized to the MGV of Lectin. (K) Western blot for KIR4.1 showing that overall protein levels are not reduced in the brain of 2-month-old *Dag1^C667F/C667F^* mice compared to heterozygous or wild-type mice. (C,F,G,I) Scale bars: 20 μm. (D,H,J) MGV analysis: n=6 Ctrl and 6 Hom animals, MGV was analyzed in 3 images per animal and included in the statistical analysis. Error bars: SEM. Statistical analysis was performed by Student’s t-test. *p<0.05 **p<0.01. (E, K) Quantification of protein bands is shown as average (n=3 mice for each genotype) and normalized to Tubulin levels (Tubulin levels shown in E also correspond to the samples blotted in K).

### The molecular composition of the glia-vascular unit is impaired in the retina of *Dag1^C667F/C667F^ mice*

Next, we examined the blood-retinal barrier at several postnatal stages (**Figure 7A**). The blood-retinal barrier is formed by the PVE of Müller glia cells at blood vessels in the deep and intermediate plexus and by PVE of astrocytes at blood vessels in the superficial plexus (**Figure 7B**) (Nicchia *et al*, 2016). The glial components of the blood-retinal barrier also contain DGC-AQP4-KIR4.1 complexes (Haenggi & Fritschy, 2006) and immunostaining for the DG core protein showed its localization around blood vessels in retinas of control animals. In the retinas of homozygous animals, the DG core protein was no longer detected around blood vessels (**Figure 7C**). Immunostaining for glycosylated α-DG or β-DG could not be performed because the available antibodies did not work on paraffin sections of the retina. Laminin, which is part of the basement membrane around blood vessels in the retina (Gnanaguru *et al*, 2013), was not altered in homozygous mutant mice (**Figure 7D**). Immunohistological analysis for AQP4 and KIR4.1 showed that in the retina of *Dag1^C667F/C667F^* mice, AQP4 was absent from the PVE at blood vessels in the superficial plexus, whereas KIR4.1 localization appeared to be normal or was only slightly reduced. In contrast, AQP4 expression persisted, albeit at reduced levels, in the PVE at blood vessels in the deep and intermediate plexus of homozygous mutant mice, while KIR4.1 was severely reduced (**Figures 7E,F**). α-SNT expression appears to be diffuse in the retina and is not restricted to the blood-retinal barrier (Enger *et al*, 2012). No difference in α-SNT localization was detected between control and mutant animals at 2- or 6-months of age (data not shown). These data demonstrate that the mutation in β-DG has distinct effects on DGC-AQP4-KIR4.1 complex formation in the PVE formed by astrocytes versus Müller glia cells, consistent with previous data suggesting that the DGC-AQP4-KIR4.1 complex differs between these two glial cell types in the retina (Enger *et al*, 2012; Nicchia *et al*, 2016).

**Figure 7.**
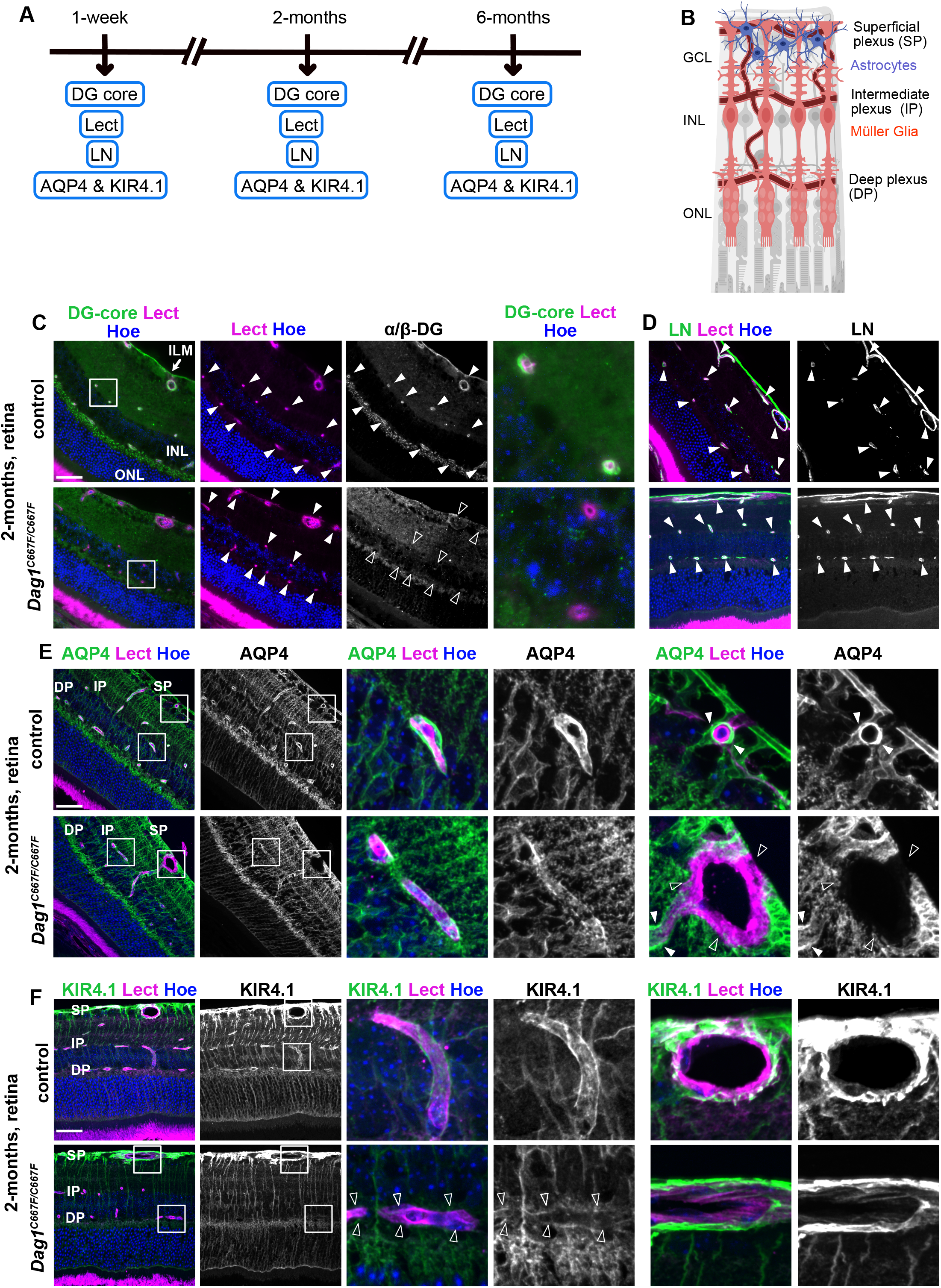
The molecular composition of the glia-vascular unit is disrupted in the retina of *Dag1^C667F/C667F^ mice*. (A) Timeline summarizing which experiments were performed at which postnatal stages. (B) Schematic of the retina showing the vasculature and the location of astrocytes and Müller glia cells. Schematic drawing generated with Biorender. (C) Immunostaining for the DG core protein (α/β-DG) in combination with Lectin (Lect) and Hoechst (Hoe) in the retina of 2-month-old mice. The DG core protein is not detectable around blood vessels in the *Dag1^C667F/C667F^* mice. Right panel: higher magnification of the boxed area in the left panels. Filled arrowheads indicate blood vessels (Lect Hoe) or α/β-DG surrounding these blood vessels (DG core). Empty arrowheads indicate absence of DG around blood vessels. (D) Immunostaining for laminin (LN) in combination with Lect and Hoe in the retina of 2-month-old mice. LN is retained at blood vessels in the homozygous mutant mice. (E) Immunostaining for AQP4 in combination with Lect and Hoe shows that AQP4 is absent around blood vessels in the superficial plexus but maintained in the intermediate and deep plexus in the retina of 2-month-old *Dag1^C667F/C667F^*mice. (F) Immunostaining for KIR4.1 in combination with Lectin and Hoechst shows that KIR4.1 is reduced around blood vessels in the intermediate (empty arrowheads) and deep plexus but maintained in the superficial plexus in the retina of 2-month-old *Dag1^C667F/C667F^* mice. (E, F) Boxes in the left panel indicate higher magnifications shown on the right. Scale bars: 40 μm.

## Discussion

### DG expression levels and localization in the *Dag1^C667F/C667F^* mice and in human patients

Our results show that DG protein levels are reduced in skeletal muscle and brain of the *Dag1^C667F/C667F^* mice. However, the glycosylation of α-DG is not affected and the localization of α/β-DG at the sarcolemma of muscle fibers is not overtly altered in the homozygous mutant mice. This observation contrasts with the reports on human patients in whom glycosylated α-DG was no longer detectable at the sarcolemma (Geis *et al*, 2013). These data suggest that the mutation may affect the processing of the DG protein differently in humans and mice. This is supported by our observations in vitro, as in a human kidney cell line (293-EBNA) transfected with the mutant DG, the folding and structure of the α/β maturation interface was severely altered (Signorino *et al*, 2018). While these in vitro results are inconclusive because they are based on overexpression of mutated murine DG in a non-muscle human cell type, they still support a potential difference in protein processing between humans and mice. A more pronounced reduction of DG at the cell membrane of human cells may be an important reason why an early-onset, mild dystrophic phenotype is observed in patients but not in the mouse model.

Despite these differences between mouse and human, the reduction in DG levels that we have observed in the mouse model allows us to make some considerations. Indeed, in transfected cell lines, the mutation does not seem to completely inhibit the maturation of the DG precursor into its two different subunits (Signorino *et al*, 2018). It also seems likely that some fraction of the unprocessed α/β-DG precursor would be destroyed or form aggregates (as indicated by the clustering pattern of β-DG that we observed in immunofluorescence at the PVE), which we are unable to transfer onto gels and blotting membranes. It is noteworthy that the C667F mutation leaves C711, which is thought to form an intramolecular disulfide bridge with C677 in the ectodomain of native β-DG (Deyst *et al*, 1995; Sciandra *et al*, 2012; Watanabe *et al*, 2007). This is consistent with the pattern of small-angle X-ray scattering (SAXS) and proteolysis data on recombinant β-DG’s ectodomain molecules, suggesting the presence of a tightly packed conformation (Signorino *et al*, 2018).

These results are relevant in several respects. First, our data strongly suggest the presence of some sort of compensatory mechanism in the mouse *in vivo* that allows for the correct processing of at least part of the DG precursor. Second, the presence of a reduced amount of properly processed, glycosylated and trafficked DG appears to be sufficient for adequate DG axis function in the development and maintenance of muscle, brain, and eye. Third, while we cannot determine whether the observed phenotypes are solely due to the reduced DG levels or whether an altered structure and interaction capacity of β-DG contributes to the phenotype, it is important to note, that the DGC appears to remain intact in the skeletal muscle and is not functionally impaired in the radial glia scaffold during development but is disrupted at the blood-brain and blood-retina barrier. These results suggest tissue-, cell- and maybe even stage-specific consequences of the C667F mutation on DG function.

### Late-onset signs of muscle pathology and skeletal muscle dysfunction in *Dag1^C667F/C667F^ mice*

Patients with the C669F mutation in *DAG1* have generalized muscular hypotonia and signs of muscular dystrophy (e.g., muscle fibers with moderate size variability) from early childhood (Geis *et al*, 2013). In the mouse model, some *Dag1^C667F/C667F^* male mice (6-months and older) show late onset signs of dystrophic muscle (increased percentage of fibers with central nuclei compared to control mice). This phenotype is not fully penetrant as it is not observed in all homozygous mutant male mice and is absent in female *Dag1^C667F/C667F^* mice. Dystrophic muscle typically exhibits greater variability in muscle fiber diameter than wildtype muscle. In our mouse model, histograms of the minimal Feret’s diameter showed a wider spread of values in male and female homozygous mutant mice compared to controls. However, VC did not show a significant difference between *Dag1^C667F/C667F^*mice and controls. Despite this mild muscle phenotype, 1-year-old male and female *Dag1^C667F/C667F^* mice were impaired in their performance on an activity wheel. In particular, the running speed of *Dag1^C667F/C667F^*mice was reduced compared to controls, suggesting that muscle function was impaired, possibly as a consequence of the observed reduction in DG protein levels. In contrast, neither motivation to run nor coordination seemed to be affected in the mutant mice.

### Disruption of the DGC and associated proteins in PVE at the blood-brain and blood-retinal barrier of *Dag1^C667F/C667F^* mice

We show that the DGC, AQP4 and KIR4.1 localization in PVE is disrupted in homozygous *Dag1^C667F/C667F^* mice. AQP4 is known to be critical for water homeostasis in the brain and retina. AQP4-null mice have altered barrier function of PVE in the brain and the deep plexus of the retina, decreased blood-brain water uptake and are more prone to develop a hydrocephalus (Haj-Yasein *et al*, 2011; Verkman *et al*, 2006; Nicchia *et al*, 2016). KIR4.1 co-localizes with AQP4 in the PVE. It has been proposed that the coupled transport of both water and K^+^ contributes to astrocytic volume changes following neuronal activity (Paz & Gulias-Cañizo, 2022).

It has previously been shown that the DGC is associated with AQP4 and plays an important role in the localization of AQP4 and KIR4.1 to PVE (Menezes *et al*, 2014; Cherkaoui *et al*, 2021; Nicchia *et al*, 2008; Sato *et al*, 2018; Enger *et al*, 2012; Sene *et al*, 2009; Rurak *et al*, 2007). For example, conditional inactivation of *Dag1* in the CNS results in greatly reduced localization of AQP4 at the PVE and an impairment of the blood-brain barrier (Menezes *et al*, 2014). Loss of Dp71 in mice, a major dystrophin gene product in the adult brain and a direct interacting partner of DG, results in a strong reduction of AQP4 in PVE and a complete absence of α-SNT in the brain and an impaired blood-retinal barrier. In addition, β-DG localization in the PVE is reduced by about half in the brain (Cherkaoui *et al*, 2021; Fujimoto *et al*, 2020; Sene *et al*, 2009). Dp71 interacts with both AQP4 and KIR4.1 in cortical astrocytes and in Müller glia cells suggesting that the DGC is essential for the localization of both proteins in the PVE (Fujimoto *et al*, 2023; Sene *et al*, 2009).

Taken together, the severe disruption of the molecular complexes at the PVE in brain and retina of *Dag1^C667F/C667F^* indicates that β-DG function is essentially lost in this highly specialized cell compartment of glia cells. This is in sharp contrast to the unimpaired development of the brain and retina and the mild muscle phenotype in the mutant mice suggesting that β-DG function is partially or fully maintained in these tissues or developmental stages despite the mutation. Importantly, the function of β-DG at the blood-brain barrier may not be restricted to the PVE, as it is also expressed in endothelia cells and pericytes (our data and (Zaccaria *et al*, 2001). Finally, the DGC has been shown to have several functions at synapses (Jahncke & Wright, 2022). Whether the C667F *Dag1* mutation also affects β-DG function at synaptic contacts in the central nervous system remains to be investigated.

### Primary dystroglycanopathies and mouse models for muscular dystrophies

While numerous cases of secondary dystroglycanopathies have been described, few patients affected by primary dystroglycanopathies have been identified so far (Brancaccio, 2019). Besides the C669F mutation in *DAG1* found by Geis and colleagues (Geis et al., 2013), which inspired the present mouse model, only three other mutations have been characterized in more detail. The observed phenotypes ranged from an early-onset form of limb-girdle muscular dystrophy (LGMD) with cognitive impairment first detected in a 3 year-old child (T192M mutation) (Dinçer *et al*, 2003; Hara *et al*, 2011) to a mild case of muscular dystrophy with hyperCKemia in a compound heterozygous 7-year-old patient (V74I, D111N) (Dong *et al*, 2015) and a late-onset form of LGMD identified in a 64-year-old man (R776C) (Dai *et al*, 2019). As far as mouse models of primary dystroglycanopathies are concerned, the only one available so far carries the missense mutation T190M (orthologous to the human T192M), which is located within the N-terminal domain of α-DG (Bozic *et al*, 2004) and results in partially impaired glycosylation of α-DG. While this mouse model shows neuromuscular abnormalities, histological signs of muscular dystrophy (i.e., centrally located nuclei in skeletal muscle fibers) have only been reported in 21-week- and 1-year-old mice (Hara et al., 2011), suggesting that the mouse model shows a late-onset muscular dystrophy phenotype, whereas the patient had an early onset of the disease. The effect of the mutation on the brain has not been investigated in the mouse model. Thus, both the T190M and the C667F mouse models show a discrepancy in the onset and severity of the phenotype compared to the human patients. Importantly, this discrepancy between mouse and human phenotypes it is not limited to these two models of primary dystroglycanopathies: even the paradigmatic *mdx* (dystrophin-less) mice, which are viable and have an almost normal lifespan, show a much milder phenotype than human Duchenne patients (Putten *et al*, 2020).

### Implications of the *Dag1^C667F/C667F^* mouse model for the treatment of rare dystroglycanopathies

It should be noted that although the phenotype of the *Dag1^C667F/C667F^*mouse model is less severe than the disease symptoms in human patients, the late onset histopathology in skeletal muscle, along with abnormalities in the PVE in the brain and retina, may well be characteristic of the mild end of the MEB disease spectrum, making this mouse model a valuable tool to study certain mechanistic aspects of this primary dystroglycanopathy. In particular, the patients with the *DAG1* C669F mutation present with megalencephalic leukoencephalopathy with subcortical cysts (MLC) (Geis *et al*, 2013). In the vast majority of patients with MLC, the phenotype is associated with pathogenic variants in *MLC1* or *GLIALCAM*. Recently, in MLC patients without mutations in these two genes, a pathogenic variant of *AQP4* was discovered that disrupts the membrane localization of AQP4 (Passchier *et al*, 2023). Thus, our finding that AQP4 localization to PVE is disrupted in *Dag1^C667F/C667F^*mice may provide a first insight into the potential mechanisms underlying the MLC in the patients with the C669F mutation in *DAG1*. Thus, we anticipate that the availability of the *Dag1^C667F/C667F^* mouse model for further molecular or pharmacological studies may have a biomedical impact, since mutations in DG, or in the enzymes responsible for its maturation in the so-called secondary dystroglycanopathies, are involved in numerous rare neuromuscular disorders that severely affect the lives of thousands of patients and families worldwide. An example of the impact that basic biochemical and cell biological research can have on the development of therapeutic strategies is the work on the DG-agrin interaction (Gesemann *et al*, 1996, 1998). Such work eventually led to the development of innovative therapeutic strategies that were developed and tested in mouse models (Moll *et al*, 2001; Meinen *et al*, 2007). Another potentially directly relevant therapeutic approach in the context of the PVE phenotype in our mouse model is the observation that AQP4 polarization in PVE is altered in a rat retinal injury model. This deficit can be rescued by treatment with bumetanide, an NKCC1 (Na+-K+-2Cl– cotransporter 1) inhibitor that downregulates AQP4 by interfering with the metalloproteinase 9 (MMP9)-mediated cleavage of β-DG (Chen *et al*, 2022). Last but not least, a better knowledge of the DG core protein can guide the design of better antibodies, especially for diagnostic purposes (Humphrey *et al*, 2014; Fortunato *et al*, 2014).

## Acknowledgment

This work was supported by the German Research Foundation (417960915 to S.B; SCHO 820/6-1 to S.S.), the SFB 1089 (to S.B., S.S.), the Bonner Promotionskolleg “NeuroImmunology” (BonnNi; 2020-S1-06 to R.L.T.) funded by the Else Kröner Fresenius Stiftung, and the French AFM-Telethon Pr. 20009 (“Establishing new models for primary dystroglycanopathies” to A.B.). We thank Karin Kappes-Horn for her guidance and support with the histological preparation and analysis of muscle tissue, Janine Marx in guidance with the histological preparation and analysis of eyes and retina, Nesrine Melliti for support with activity wheel analysis, the UKB microscopy core facility for support with imaging; Khondker Ushna Sameen Islam and Norisa Meli for support with mouse work.

